# Multiscale Modeling of MT1-MMP-Mediated Cell Migration: Destabilization of Cell-Matrix Adhesion

**DOI:** 10.1101/2022.10.12.511909

**Authors:** V. Andasari, M. Zaman

**Affiliations:** Department of Electrical Engineering, University of Indonesia, Depok 12455, Indonesia; Department of Biomedical Engineering, Boston University, Boston, MA 02215, USA

## Abstract

One of several ways MT1-MMP promotes cell migration is by modifying cell adhesion properties. MT1-MMP directly processes cell adhesion properties by shedding cell transmembrane receptors that attach cells to the extracellular matrix (ECM). The shedding leads to the destabilization and disassembly of firm cell-matrix adhesion that holds cells in their stationary position, prompting cells to migrate. In this paper, we present a multiscale mathematical model of single cell migration driven by MT1-MMP destabilization of cell-matrix adhesion. The dynamics of MT1-MMP are modeled using a system of differential equations that are integrated with the Cellular Potts Model (CPM) for a combined modeling at the intracellular and cellular scale, respectively. The CPM is extended to include a local feedback mechanism from MT1-MMP on the membrane that enhances cell membrane fluctuations, resulting in actively migrating cells. The results of computational simulation show that MT1-MMP is capable of destabilizing strong cell-matrix adhesion and stimulating cell migration, and at the same time, also producing cell polarization and motile cell morphology.

## 1 Introduction

Cell migration is a complex and highly organized cellular activity that plays a central role in many physiological and pathological conditions. To migrate, cells respond to extracellular stimuli, such as substrate properties, cytokines, and growth factors, by activating various classes of protease. There are several proteases that are known to be involved in cell migration. In particular, matrix metalloproteinases (MMPs) have been extensively studied for their potent impact on cell migration. MMPs consist of a family of zinc-dependent enzymes. To date, there are at least 23 different types of MMPs expressed in human tissue [1]. Of these, membrane-tethered membrane type 1-matrix metalloproteinase (MT1-MMP) has emerged as a key member. Specifically in cancer studies, MT1-MMP is closely associated with tumor invasiveness and malignancy [2]. MT1-MMP is also implicated in other diseases and abnormalities such as dwarfism, osteopenia, progressive fibrosis, arthritis, impaired angiogenesis, and pulmonary hyperplasia [3, 4, 5].

MT1-MMP, also known as matrix metalloproteinase-14 (MMP-14), was first identified in 1994 by Sato et al. as an activator of proenzyme matrix metalloproteinase 2 (proMMP-2) on the surface of invasive lung carcinoma cells [6]. Since its discovery, MT1-MMP has been extensively researched experimentally and theoretically, and is now one of the best characterized MMPs. MT1-MMP is synthesized as an inactive proenzyme with a multidomain structure. Each MT1-MMP domain plays a prominent role in extracellular matrix (ECM) degradation and cell migration. The MT1-MMP domain consists of a signal peptide, a propeptide, a catalytic domain (CAT) with a catalytic zinc atom (Zn) for proteolytic activity, a hinge region (L1, linker-1), a hemopexin-like (Hpx) domain that interacts with cell surface molecules such as CD44 and homodimer formation, a stalk (L2, linker-2) region, a type-1 transmembrane domain (TM), and a cytoplasmic tail (CT) for interactions with *β*_1_ or *α_v_β*_3_ integrins [7, 8, 9, 10, 11]. During translation, the signal peptide is removed to generate the proMT1-MMP zymogen [12, 13]. The proMT1-MMP zymogen is activated into its mature form, MT1-MMP, by removing the propeptide domain [14, 15, 16] by several members of the pro-protein convertase (PC) family such as furin, PC5/6, PC7, PACE4 [14, 17, 13], all of which are ubiquitously expressed [18]. The activation of MT1-MMP occurs intracellularly during secretion in the Golgi [19, 13].

As a membrane-anchored enzyme, active MT1-MMP is localized and anchored to the cell membrane via the transmembrane domain followed by the cytoplasmic tail [20, 21]. Since MT1-MMP is localized to the cell membrane, its activity is confined to the immediate microenvironment surrounding cells or the pericellular space. Once expressed on the cell membrane, MT1-MMP is anchored to surface structures. In non-neoplastic cells, the localization of MT1-MMP on the cell membrane is located at lamellipodia (on a 2-dimensional or 2D substrate) or filopodia (in a 3-dimensional or 3D environment) and in cancer cells at invadopodia [22] via the transmembrane domain [23]. All these protrusive membrane structures are particularly rich in actin dynamics [24].

The main biological function of MT1-MMP is to promote cell migration using various means, such as (i) by degrading components of the pericellular ECM to create a migration path, (ii) by modifying cell adhesion properties by shedding cell adhesion molecules to increase cell motility and attenuating integrin clustering to induce cell migratory signaling, and (iii) by altering cellular metabolism [25, 26, 10, 27]. Regulation of MT1-MMP as a proteinase in ECM proteolysis is well established, and the mechanisms of MT1-MMP-based proteolysis are fairly well understood, where MT1-MMP can directly [25] or indirectly [6, 28, 29] degrade ECM components in order to create a path for cell migration.

In addition to its role in extracellular proteolysis, MT1-MMP also stimulates cell migration by directly processing cell transmembrane receptors that attach cells to the ECM. It is achieved by shedding receptors from the cell surface as well as reducing the clustering of cell transmembrane receptors. These activities destabilize and disassemble cell-matrix adhesions, leading to upregulation of adhesion turnover and inducing cell migration [30, 26]. MT1-MMP has been shown to shed CD44H in various cancer cell lines, such as human pancreatic carcinoma, breast carcinoma, and osteosarcoma [30, 31, 32]. MT1-MMP forms a complex with CD44H via its Hpx domain that binds directly to the stem region of CD44H [33, 34]. CD44H (or hematopoietic CD44) is a cell surface adhesion molecule that participates in cell-cell and cell-matrix adhesion and is highly expressed in many types of cancer cells [35]. CD44 is a family of cell surface glycoproteins, where CD44H is one of its isoforms [36]. For its function in cell-matrix adhesion, CD44 binds to extracellular matrix ligands, including hyaluronan or hyaluronic acid (HA), osteopontin, collagen, fibronectin, laminin, and chondroitin sulfate, with HA being the main ligand [35, 37]. Other transmembrane receptors that are targets of shedding by MT1-MMP to stimulate cell migration and cancer malignancy include, among others, *α_v_*-integrin [38, 39], syndecan 1 [40], ICAM-1 [41], LRP1 [42], mucin16/CA-125 in ovarian cancer cells [43], and death receptor-6 (CR6) [44].

MT1-MMP has been reported to shed CD44 from the cell surface by proteolytically cleaving it in its stem region and releasing a soluble fragment of CD44H in the ECM [30, 33, 34, 31, 32]. Since the cytoplasmic domain of CD44 binds to the filamentous action (F-actin) via Ezrin/Radixin/Moesin (ERM) proteins [32], Shedding of CD44 causes the cell to detach from its fibrillar adhesion structures and migrate through the matrix [30]. As we have previously studied, only medium-sized adhesive structures, such as nascent adhesions and focal complexes, that enable cell migration whereas large adhesive structures, such as fibrillar adhesion, act as a stable scaffolding that prevents cells from migrating [45]. Several other molecules have been shown to mediate CD44 shedding by MT1-MMP, such as TGF-*β* [46], mDia1 [47], EGFR [48], and KIF5B and KIF3A/KIF3B [49]. Although these studies show that CD44 shedding by MT1-MMP occurred at the front edge, shedding could also occur at the rear end of invasive tumor cells for cell retraction during migration [50]. The interactions of MT1-MMP and CD44 were found to be involved in the motility of various types of cancer, such as pancreatic cancer [51], ovarian cancer [52], and breast cancer [46, 48, 31, 47].

Although mathematical models on the role of MT1-MMP in ECM proteolysis exist, such as a model of the biochemical network of MT1-MMP, MMP-2, and TIMP-2 [53], a model of the secretion of diffusible MMP-2 after binding to MT1-MMP and TIMP-2 out of a single cylindrical cell surrounded by a 3D collagen type I matrix [54], the importance of rapid MT1-MMP turnover for proteolysis [55], an analysis of the formation of ternary complex MT1-MMP/MMP-2/TIMP-2 [56], MT1-MMP-related invadopodia modeling [57, 58, 59] as well as the implications of MT1-MMP in cancer invasion [60, 61, 62], to our knowledge, there have not been developed mathematical models on the role of MT1-MMP in modifying cell adhesion property by destabilizing and disassembling cell adhesivity to the ECM for cell migration. Therefore, in this paper, we present a multiscale individual cell-based model of cell migration that is driven by the destabilization of MT1-MMP and the disassembly of cell matrix adhesion. In our model, a single cell is assumed to have two compartments: an intracellular cytoplasmic compartment and a membrane compartment. We assume that the activation of proMT1-MMP into active MT1-MMP occurs in the cytoplasmic compartment. The dynamics of proMT1-MMP and active MT1-MMP are modeled using ordinary differential equations (ODEs). Active MT1-MMP is then transported to the membrane compartment for co-localization with cell adhesion receptors, which are assumed to take place at invasive structures. Membrane MT1-MMP is also internalized through a feedback mechanism. The schematic diagram of the process is shown in Fig. 1. ODEs that model intracellular MT1-MMP dynamics are incorporated into cellular modeling using the lattice-based Cellular Potts Model (CPM). Our simulation results show that incorporating MT1-MMP dynamics enhances cell migration, producing motile cells with polarization and motile morphology.

**Figure 1:**
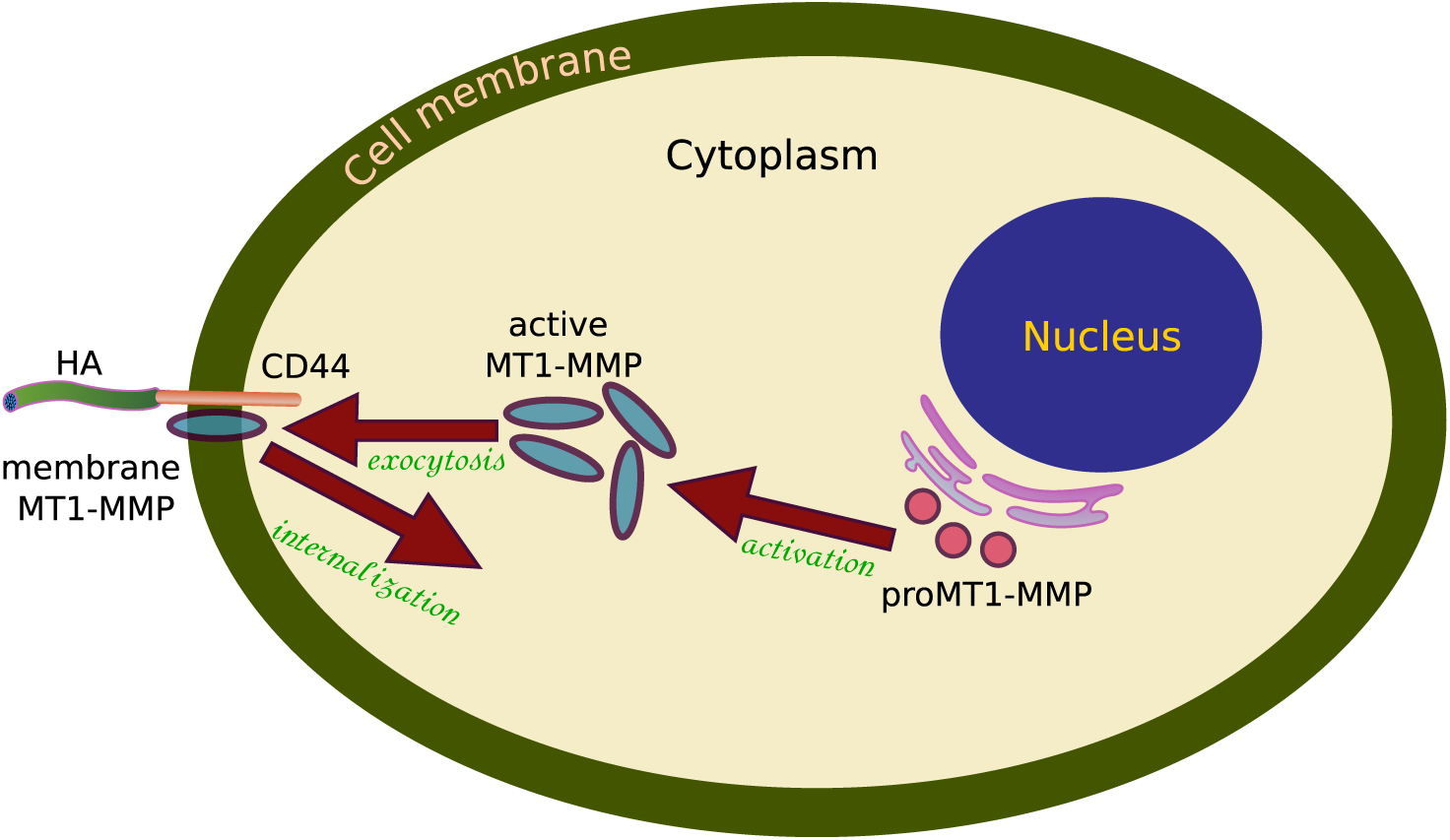
Schematic diagram showing the compartmental model describing the interactions between intracellular MT1-MMP (inactive and active) concentrations and membrane MT1-MMP for cell migration.

## 2 The Mathematical Modeling & Computational Framework

We implemented the dynamics of intracellular and membrane of MT1-MMP in single cells using the CPM, also known as the Glazier-Graner-Hogeweg (GGH) model [63, 64, 65, 66]. The dynamics of single cells in the CPM are described by an effective energy formalism known as the Hamiltonian and denoted by 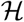. The Hamiltonian in our model contains the terms for adhesion energy between cell-cell and cell-matrix, volume and surface constraints, membrane constraint, as well as chemotaxis constraint for simulations with directed migration. All these terms are formulated as

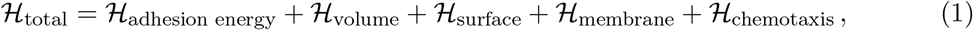

where

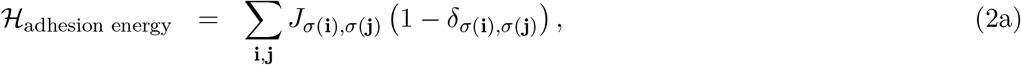

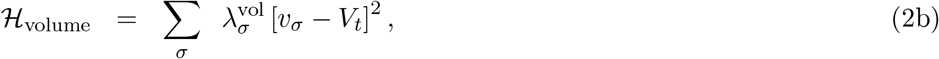

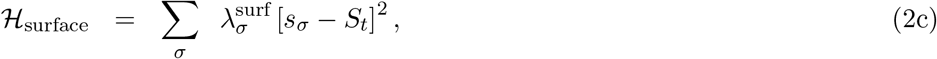

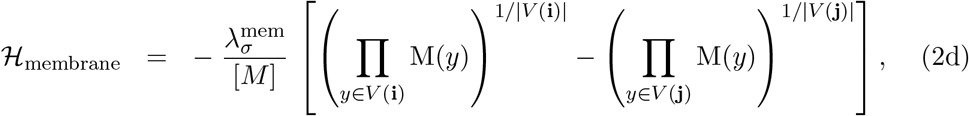

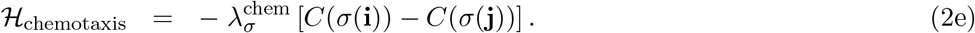

The first term, Eq. (2a), represents variations in energy due to adhesion between cells of different types (including the medium or ECM) with *J*_*σ*(**i**)*σ*(**i**)_, denotes a boundary energy between cells *σ*(**i**) and *σ*(**j**), and *δ*_*σ*(**i**),*σ*(**j**)_ denotes the Kronecker delta function, given by

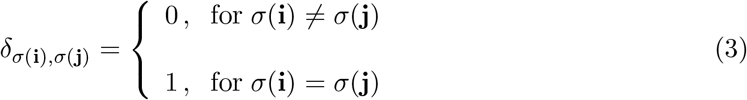

where the factor (1 – *δ*_*σ*(**i**),*σ*(**j**)_) ensures that only pixels belonging to different cells are counted. The summation runs over all neighboring pairs of lattice sites **i** and **j**. In the second term, Eq. (2b), which represents a volume constraint, 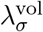 denotes the volume elasticity parameter, *v_σ_* the cell’s actual volume, and *V_t_* the cell’s target volume. The summation over *σ* runs over all cells in the lattice. Likewise for the third term, Eq. (2c), which represents a surface constraint, 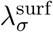 denotes the area elasticity parameter, *s_σ_* the cell’s actual area/perimeter, and *S_t_* the cell’s target area. Following [45], *S_t_* is calculated using circularity which measures cell morphology, given by

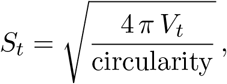

where, if the value of circularity *S_t_* is close to 1.0, it leads to a perfect circle of a cell shape, whereas if the value of *S_t_* approaches 0, the cell is increasingly elongated.

The fourth term, Eq. (2d), is adopted from [67] and we refer to here as the membrane model. This term represents the energy due to a membrane constraint, where 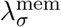 denotes the maximum contribution of the membrane model to the system energy, [*M*] the concentration of membrane MT1-MMP, and (⋂ *M*(*y*))^1/|*V*(*x*)|^ the geometric mean of the activity values in the neighborhood of *x* with *V*(*x*) the direct Moore neighbors of *x*. The last term, Eq. (2e), represents interactions of cells with an external chemical gradient *C* with 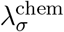 denotes the cell’s chemotactic response parameter. A cell will move up-gradient if 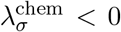 or down-gradient if 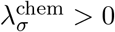 [65].

The CPM has been widely used to model cell behaviors and interactions for prediction at the cellular- and tissue-scale. The model has also been combined with other mathematical techniques to simulate various and more complex cell behaviors. One of our previous CPM-based models was the first that successfully integrated the intracellular dynamics of proteins involved in cell-cell adhesion with individual cell dynamics to model cancer growth and invasion [68]. In the model, the intracellular equations were described using the Systems Biology Markup Language (SBML) which were then translated into ODEs coupled with the CPM.

The majority of simulations we performed for this paper were in 2D, whereas 3D simulations were performed only for a chemotaxis model. The following modeling mechanism applies for 2D simulations, while 3D modeling is explained in the Result section for chemotaxis. We developed a CPM computational framework for multiscale modeling without using the SBML. The framework directly integrates the intracellular dynamics formulated in a system of ODEs using an ODE solver and couples the results with the CPM’s cellular dynamics. The ODE solutions are integrated with the cellular dynamics after each index-copy attempt of the CPM; hence, the intracellular dynamics affect the next index-copy attempt. Our framework is quite flexible, and implementing intracellular dynamics using SBML files can still be performed if preferred. We can simulate as many ODEs for intracellular dynamics and the ODEs can be either stiff or nonstiff problems.

We made several assumptions in the formulation of our MT1-MMP model:

- The process starts with enzyme synthesis. Inactive proMT1-MMP, denoted by *P*, is synthesized at a constant rate *k_s_*.
- The processing (or activation) of proMT1-MMP into active (or mature) MT1-MMP occurs intracellularly. In the activation process there are no bindings with any activator because activation by furin means cleavage of 61 kDa proMT1-MMP into 39 kDA and 22 kDA fragments [14, 17]. The state of inactive proMT1-MMP is changed into active MT1-MMP, denoted by *A*, with a rate of *k*_1_.
- Active MT1-MMP is then transported to the cell membrane, becoming membrane-bound MT1-MMP denoted by *M* which is employed by cells to destabilize strong cell-matrix adhesion.

With these assumptions, we propose the dynamics of proMT1-MMP as follows. Disregarding gene transcription process, inactive proMT1-MMP concentration, denoted by [*P*], is synthesized at a constant rate *k_s_*. Its activation to MT1-MMP is formulated using a nonlinear degradation term with rate *k*_1_. The rate of change of [*P*] is given by

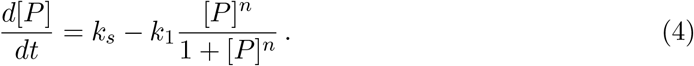

The rate of change of active MT1-MMP, whose concentration is denoted by [*A*], is modeled by linear production with rate *k*_3_ which is assumed to come from: (i) internalization (or endocytosis) with rate *k*_3_ and is in active form, (ii) nonlinear production from proMT1-MMP with rate *k*_1_, and (iii) decay with a rate *k*_2_ which is assumed to be a combination of exocytosis and destruction in lysosomes,

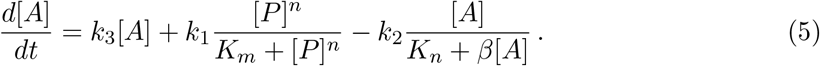

The parameter values for simulating the model equations (4)–(5) for 2D simulations are given in Table 1. These values are intended to obtain a continuous supply of active MT1-MMP. The solutions of the ODEs are shown in the third column. The parameter set has initial conditions of 5.5 concentration units for inactive proMT1-MMP and 1.0 for active MT1-MMP.

**Table 1:** Estimated parameter values for 2D MT1-MMP dynamics

| Parameter | Estimated Value |
| --- | --- |
| $k_s$ | 0.1 |
| $k_1$ | 0.16 |
| $k_2$ | 0.022 |
| $k_3$ | 0.01 |
| $n$ | 2 |
| $K_m$ | 2 |
| $K_n$ | 1 |
| $\beta$ | 0.0 |
| $\alpha$ | 2 – 3 |

After being activated, a major fraction of active MT1-MMP is transported to the cell membrane. On the cell membrane, unlike intracellular dynamics where the interactions are modeled using ODEs, membrane MT1-MMP [*M*] is assumed to be in a linear relationship with active MT1-MMP [*A*], given by

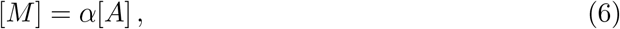

where *α* is a constant describing how much active MT1-MMP [*A*] is converted into membrane MT1-MMP [*M*] and localized at invadopodia, which are rich in actin and capable of penetrating extracellular matrix barriers. Since only moderate amounts of MT1-MMP are located at invadopodia for migration, we took the value of [*M*] ~ 16, hence 2 ≤ *α* ≤ 3. Higher amounts of MT1-MMP, such as [*M*] > 40, will generate migration behavior of a keratocyte-like cell with very wide lamellipodia, whereas very low amounts of MT1-MMP, such as [*M*] < 5 will not be able to generate enough membrane fluctuations for a cell to move its body because small numbers of [*M*] will be quickly depleted or internalized due to rapid membrane MT1-MMP turnover.

Our intracellular model does not include spatial distribution, hence we do not have equations for MT1-MMP transport from the cytoplasm to the cell membrane and MT1-MMP dispersion on the cell membrane for the 2D model. Some of MT1-MMP on the membrane is involved in ECM proteolysis and some plays a big role in disassembling of stable and strong cell-matrix adhesion by cleaving cell-matrix adhesion receptors. While MT1-MMP for ECM proteolysis forms complexes with MMP-2 and/or TIMP-2, on the other hand, MT1-MMP for destabilizing cell-matrix adhesion has been observed to co-localize with cell-matrix adhesion receptors such as integrins and CD44. In our model, this co-localization is implemented by calculating the geometric mean for the membrane MT1-MMP at the same site where cell-matrix adhesion energy of the CPM is calculated.

The CPM we implemented in this paper is based on the modified Metropolis algorithm [69], in which, at each index-copy attempt one target pixel and one source pixel are selected. If these pixels belong to the same cell ID, no copy index is performed; otherwise, the source pixel occupies the target pixel. With this occupation, if the source pixel is in a cell and the target pixel is in the medium, the cell membrane expands; if the opposite occurs, the cell membrane retracts. An index-copy attempt is the changes in the effective energy 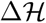 due to accepting or rejecting an attempt to move cell lattice boundaries with a probability. One index-copy attempt for each pixel in the cell lattice is called a Monte Carlo Step (MCS) [69]. By incorporating the membrane model, at each index-copy attempt the membrane constraint creates a local positive feedback that biases the copy attempt by subtracting 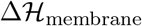 from the 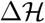 system, hence the negative sign before Eq. (2d). Then, the concentration of membrane MT1-MMP [M] is assigned as the value of the source pixel that freshly occupies the target pixel in membrane protrusion. This assignment only applies if the source pixel is in the cell. However, if the source pixel is in the medium, zero is assigned as its value.

The value of [*M*] at the freshly occupied pixel decreases by one after every MCS for memory activity of the membrane model. It resembles the internalization undergone by MT1-MMP on the cell membrane. Since some internalized MT1-MMP can be transported to lysosomes for destruction, we assume only a small fraction that is recycled to the pool of active MT1-MMP [A], hence the small value of parameter *k*_3_. Both continuous supply and internalization create rapid turnover of MT1-MMP at the cell membrane, a critical process in the regulation of cell migration.

## 3 Computational Simulation Results

We performed 2D and 3D simulations of single cell migration promoted by MT1-MMP. The code for 2D simulations was written in Cython (https://cython.org/). Cython is a superset of Python that compiles to C, hence giving our simulations a C-like performance that is many times faster than pure Python. The source code was written in Python extension functions with C-like syntax and saved in Pyrex (.pyx) files. Cython provides a Numpy module with fast access for array operations that we made use of. Some of our functions also applied C arrays for fast calculations. Visualization was performed using the Matplotlib library [70]. All 3D simulations were performed using Morpheus [71]. Additional software such as bioView3D (https://bioimage.ucsb.edu/bisque/bioview3d) and ParaView (https://www.paraview.org) were used to visualize the 3D simulation results that were saved during Morpheus simulations as `.tif` and `.vtk` files.

Random cell migration was simulated on a square lattice domain of size 100 × 100 pixels in the *x* and *y* directions. In the 2D simulations, at 0 MCS a cell with a square initial shape containing 20 × 20 pixels was placed at the center of the domain. Simulations of the up-gradient movement of a cell due to an external field, or chemotaxis, were performed in 2D and 3D. For 2D simulations, a cell was initially placed near the left end of a rectangular lattice of 200 × 70 pixels with an external static field gradually increasing toward the right end. For 3D simulations, a spherical cell of radius 10 pixels was also placed near one end of a rectangular cuboid of 200 × 100 × 100 pixels with an external static field gradually increasing toward the other end. All simulations were run up to 1000 MCS for 2D and 1500 MCS for 3D, except for simulations to measure the mean squared displacement (MSD) which were run up to 2000 MCS. And in all simulations, single cells are assumed to migrate in a homogeneous extracellular environment. All CPM parameters of Eq. (1) for 2D and 3D simulations are given in Table 2.

**Table 2:** Cellular Potts Model parameter values

| Parameter | 2D Estimated Value | 3D Estimated Value |
| --- | --- | --- |
| $T$ | 10 | 5 |
| $J_{\text{medium-medium}}$ | 0 | 0 |
| $J_{\text{cell-medium}}$ | 5, 10, 20 | 150 |
| $J_{\text{cell-cell}}$ | 0 | 0 |
| $V_t$ | 450 | 10,000 |
| $\lambda_{\sigma}^{\text{vol}}$ | 5 | 2 |
| circularity | 0.4 | 0.04 |
| $\lambda_{\sigma}^{\text{surf}}$ | 5 | 1.3 |
| $\lambda_{\sigma}^{\text{mem}}$ | 300 | 5 |
| $\lambda_{\sigma}^{\text{chem}}$ | -200 | $-2400 \times [M]_{\text{max}}$ |

Our 2D framework provides options for periodic and zero-flux boundary conditions. There are also two options for the adhesion constraint provided: adhesion energy expressed as constants and adhesion energy expressed as individual cell adhesion molecules. For our simulations, we chose to apply the periodic boundaries and adhesion energy expressed as constants, hence, constant values of *J*_medium-medium_ and *J*_cell-medium_. Interactions between pixels were implemented using the Moore neighborhood. Since our ODEs are not a stiff problem, the system of ODEs for intracellular dynamics of proMT1-MMP and active MT1-MMP in 2D simulations was solved using the explicit 4-th order Runge-Kutta method. Other explicit and/or implicit methods can also be easily applied by modifying parameters of Butcher tableau provided in the Runge-Kutta solver function. The ODEs were written in a separate function for ease of changes, either the equations or parameter values. Each MCS, the ODE solutions are directly integrated into the single cell model of CPM. Between MCS, the time interval for ODEs is discretized into 10 steps for faster calculation. All code for 2D simulations are available at https://github.com/vandasari/mt1mmp_paper.

### 3.1 The Membrane Model

We adopted the Act model developed by Niculescu et al [67] for MT1-MMP dynamics on the cell membrane. The Act model, or we refer to here as the membrane model, is based on a local feedback mechanism that can enhance membrane fluctuations resulting in a more motile cell, as shown in Fig. 2. One cell with the membrane model (bottom row figures) and one without (top row figures) were simulated with similar parameter values. Specifically, the value of cell-matrix adhesion *J*_cell-medium_ = 10 for both scenarios throughout 1000 MCS.

**Figure 2:**
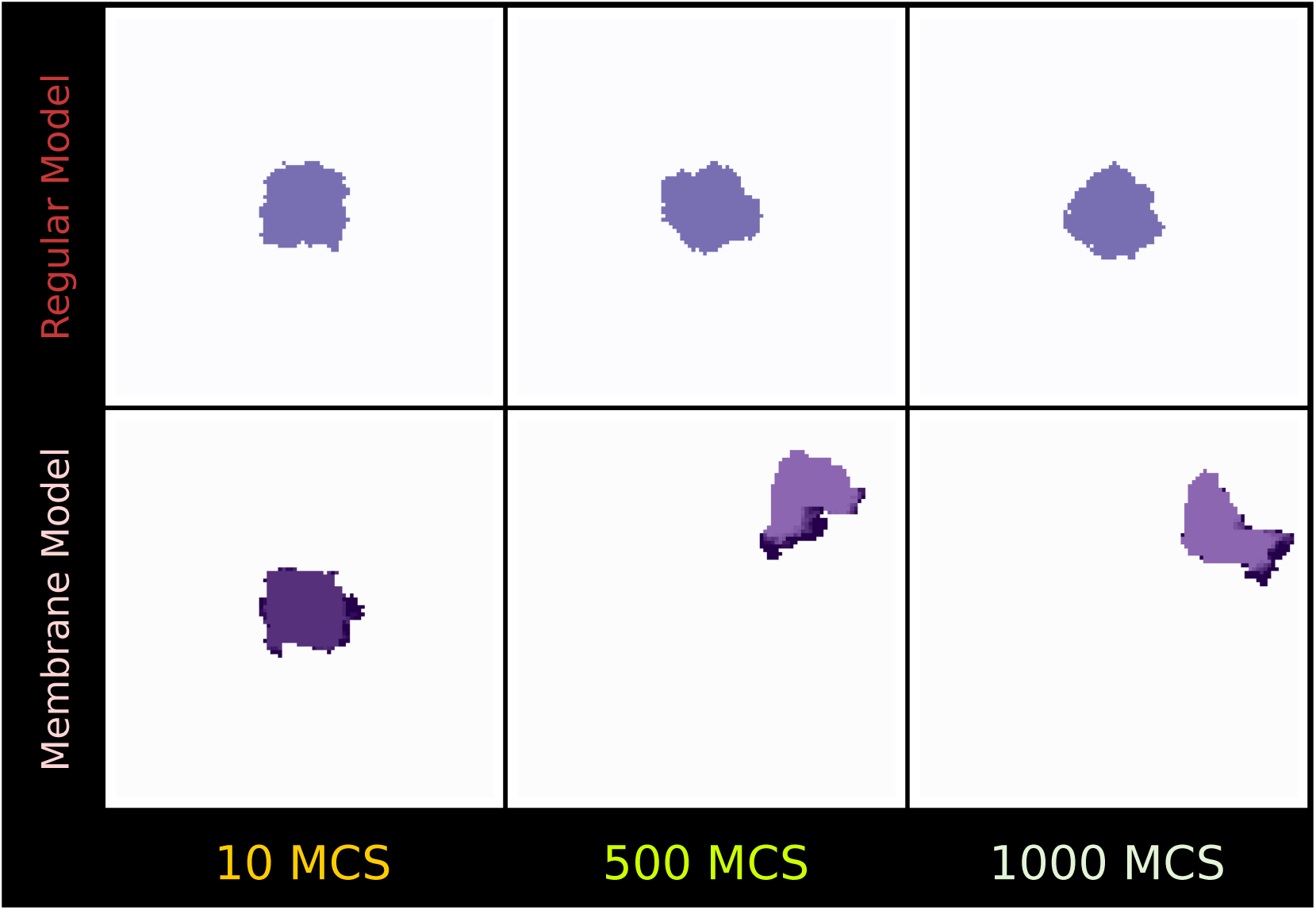
Top and bottom rows show a cell without and with the membrane model, respectively, taken at 10 MCS (left column), 500 MCS (middle column), and 1000 MCS (right column). All parameter values for simulating both cells were the same. The cell with the membrane model (bottom row) is more motile compared to the cell without the membrane model (top row). See movies 1 and 2 in the supplementary material.

The local feedback mechanism of the membrane model occurs at the sites of target pixel and source pixel, where the geometric mean of the Moore neighbors of each target and source pixels is calculated with overflow prevention. This is done by mapping the values of Moore neighbors to a logarithmic domain, summing up these logarithmic values, dividing the sum by the number of Moore neighbors, and calculating the exponents.

Rapid turnover of MT1-MMP on the membrane creates membrane expansion and retraction that have been observed in migrating cells *in vivo* and *in vitro*. In the bottom row of Fig. 2, membrane MT1-MMP [M] is visualized as a spectrum of dark purple colors in the motile cell with the membrane model. The darkest color means that the pixel is freshly assigned the value of [*M*]. The darkest pixel also indicates the front of the cell since this is where membrane protrusion occurs.

Stopping a migrating cell can be done by decreasing the concentration of active MT1-MMP [*A*], which then will reduce the concentration of membrane MT1-MMP [*M*]. This is equivalent to removing the membrane model from the cell, and then the cell will behave like the regular model, as in the first row of Fig. 2.

### 3.2 MT1-MMP Destabilizes Stable Adhesion

To simulate strong cell-matrix adhesion destabilization by MT1-MMP, we initially ran the CPM without the intracellular dynamics of MT1-MMP and the membrane model. Without the membrane model, the cell membrane slightly fluctuates causing very little random movement of the cell. Over time, the value of *J*_cell-medium_ was increased progressively until the fluctuation of the cell membrane was limited, as seen in figures with white background in the first row of Fig. 3. When the cell became immobile due to stable and strong cell-matrix adhesion with *J*_cell,medium_ = 30, the dynamics of intracellular MT1-MMP was incorporated at 400 MCS which then regulated the membrane model. A small protrusion with MT1-MMP started to appear at 450 MCS, which became bigger and after a while disassembled strong cell-matrix adhesion, triggering cell motility.

**Figure 3:**
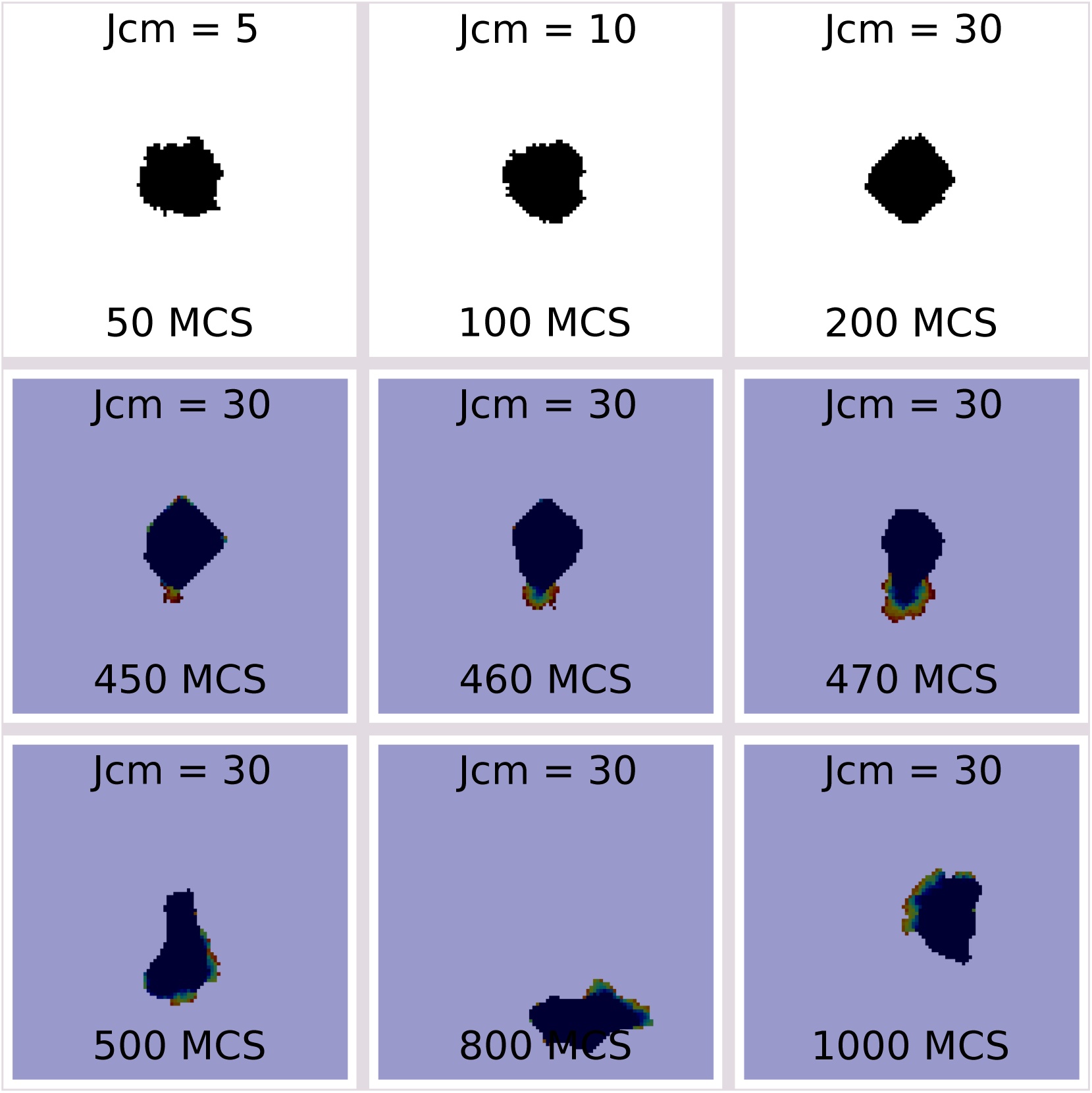
Figures showing the destabilization of strong cell-matrix adhesion by MT1-MMP. Top row with white background shows a cell without the membrane model. Over time, cell-matrix adhesion increases from *J*_cell,medium_ = 5 to *J*_cell,medium_ = 30. The intracellular dynamics of MT1-MMP that regulate the membrane model is applied at 400 MCS. Middle and bottom rows with purple background show a motile cell with the membrane model regulated by MT1-MMP. See movie 3 in the supplementary material.

The MT1-MMP dynamics also relaxed the cell membrane that was previously stiff due to strong cell-matrix adhesion. Furthermore, we observe a more diverse cell morphology, as seen at 500 MCS, 800 MCS, and 1000 MCS. Here membrane MT1-MMP [*M*] is visualized as a spectrum of the jet colormap, where dark red being the highest value of [*M*] and dark blue the lowest. The dark red pixels are where the front edge of the cell is.

### 3.3 Mean Squared Displacement and Anomalous Diffusion

We investigated the movement of a single cell with MT1-MMP dynamics on a 2D domain by plotting the mean squared displacement (MSD) over time. The definition of MSD we implemented to generate plots of MSD vs time in Fig. 4 is given by

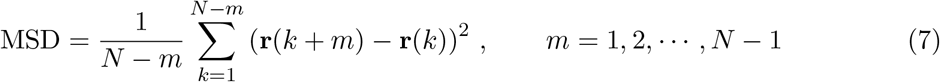

where **r**(*i*) = (*x_i_, y_i_*) is the position vector of the center of the cell and *N* is the total number of MCS. In all MSD simulations, we extended the simulation time to 2000 MCS. We performed six sets of simulation; each set represents different values of the parameter cell-matrix adhesion *J*_cell-medium_. The MSD is averaged over at least five simulations for each value of *J*_cell-medium_. To avoid problems related to boundary effects due to the finite size, we ran simulations on a larger domain of 200 × 200 pixels and only included simulations where the cell never touched or crossed the boundaries.

**Figure 4:**
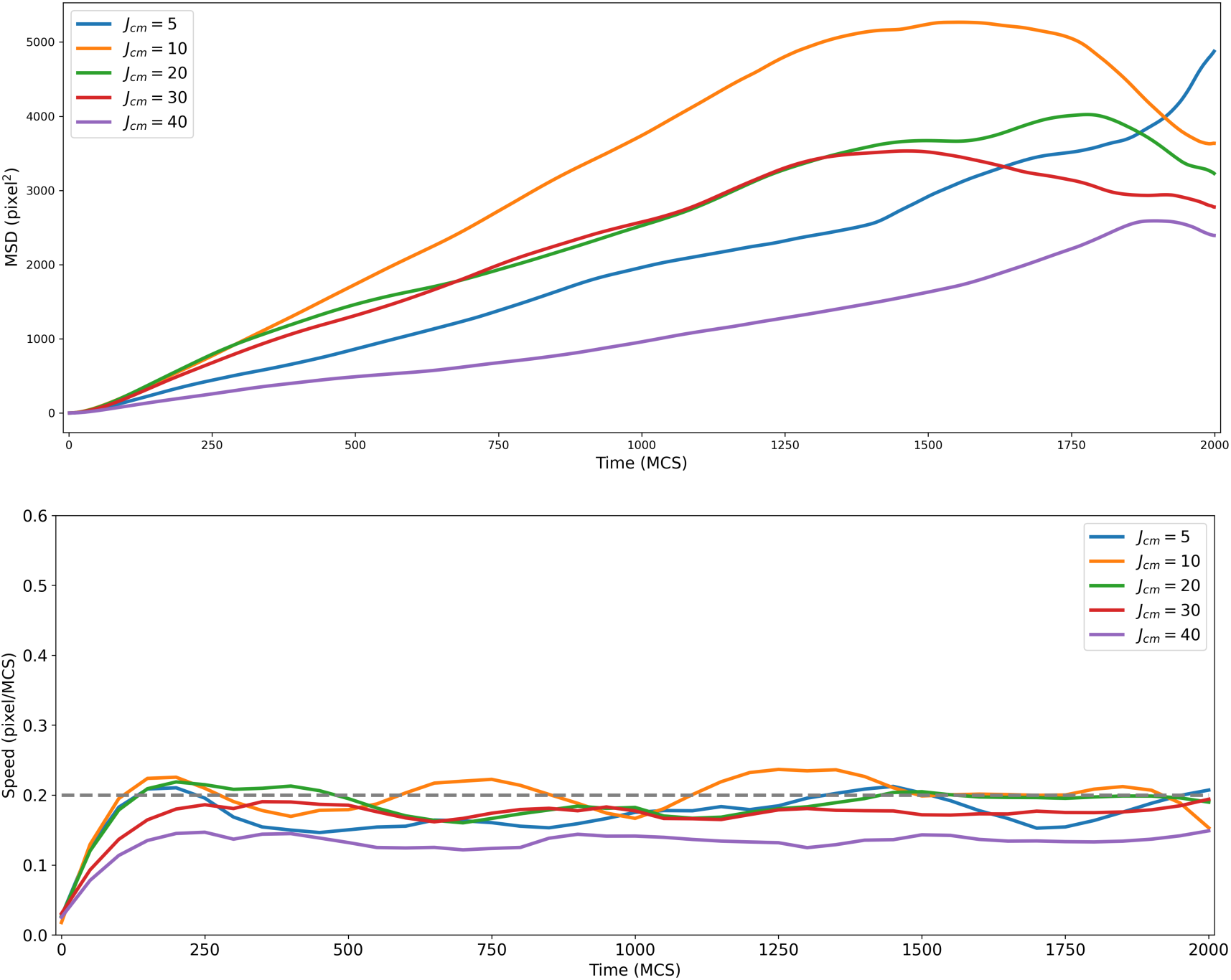
Plots of the MSD vs time (top) and the average speed vs time (bottom) for different *J*_cel-medium_ values simulated from a single cell with the membrane model.

In Fig. 4, we observe that cells with intermediate values of cell-matrix adhesion, e.g., *J*_cell-medium_ = 10, *J*_cell-medium_ = 20, and *J*_cell-medium_ = 30, they move fast, as can be seen in plots of speed vs time (MCS) at the bottom of Fig. 4. Cells having medium values of cell-matrix adhesion *J*_cell-medium_ exhibit nonlinear relationships between the MSD and time, where the curves represent what looks like an anomalous subdiffusion, *i.e.*, MS ∝ time^*θ*^ where *θ* is called the anomalous exponent. The value of *θ* and exactly equal to 1 for normal diffusion and *θ* < 1 for subdiffusion. Anomalous subdiffusion in biology has been experimentally observed for proteins and lipids on the cell membrane (2D diffusion) and for proteins and molecules in the nucleus and cytoplasm (3D diffusion) [72, 73, 74]. Anomalous diffusion in invasive cancer cells during migration and proliferation has also been the subject of study [75].

### 3.4 Chemotaxis

Directed movement of cancer cells due to extracellular chemical gradients, or also known as chemotaxis, is an important part of cancer dissemination during progression and metastasis. Chemotaxis of single cells can be categorized into amoeboid migration and mesenchymal migration. Of these two, mesenchymal migration is the mode of migration that requires the activity of MMPs to break down physical barriers and propel the cell body forward. Chemotaxis of mesenchymal cells is characterized by an elongated cell morphology with established cell polarity. We were interested in knowing whether our model could produce elongated migrating single cells as well as polarization.

We performed 2D and 3D simulations of chemotaxis, in which an external gradient was assumed to be a static chemical field whose high concentrations attracted cells to move toward. Simulations were performed on a rectangular domain for 2D and in a rectangular cuboid for 3D. The static chemical field gradually increased from one end to the other end of the long axis. At 0 MCS, we placed a cell at the end where the chemical field is low and let the cell move following the chemical gradient over time.

Applying the membrane model indeed results in chemotaxis with elongated cell morphology in both 2D and 3D, as shown in Figs. 5 and 6, respectively. For 2D, we performed four sets of simulation, each for different cell-matrix adhesion energy values. Results for *J*_cell-medium_ = 5, *J*_cell-medium_ = 10, *J*_cell-medium_ = 20, and *J*_cell-medium_ = 30 are shown in the first row, second row, third row, and fourth/bottom row figures of Fig. 5, respectively. The results shown are taken at 100 MCS (left column), 500 MCS (middle column), and 1000 MCS (right column). Like random migration in Fig. 3, the chemotactic cell establishes polarization to differentiate the front edge from the rear edge. At the front edge, the localization of MT1-MMP on the cell membrane is fairly visible at 100 MCS, as shown by the figures in the first column of Fig. 5.

**Figure 5:**
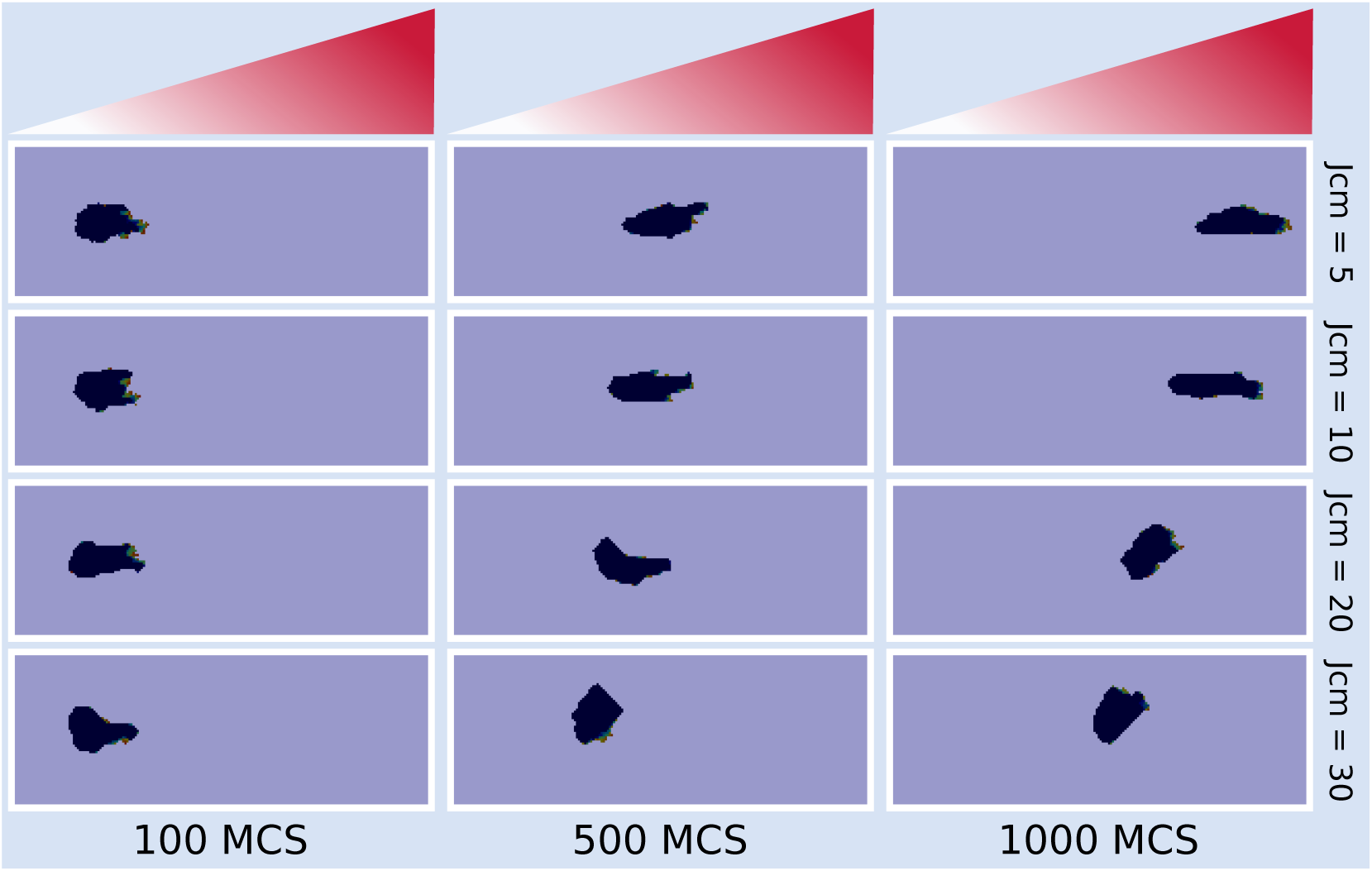
Chemotaxis of a single cell with the membrane model exhibiting elongated cell morphology, simulated with *J*_cell-medium_ = 5 (first row), *J*_cell-medium_ = 10 (second row), *J*_celll-medium_ = 20 (third row), and *J*_cell-medium_ = 30 (fourth/bottom row), each taken at 100 MCS (left column), 500 MCS (middle column), and 1000 MCS (right column).

**Figure 6:**
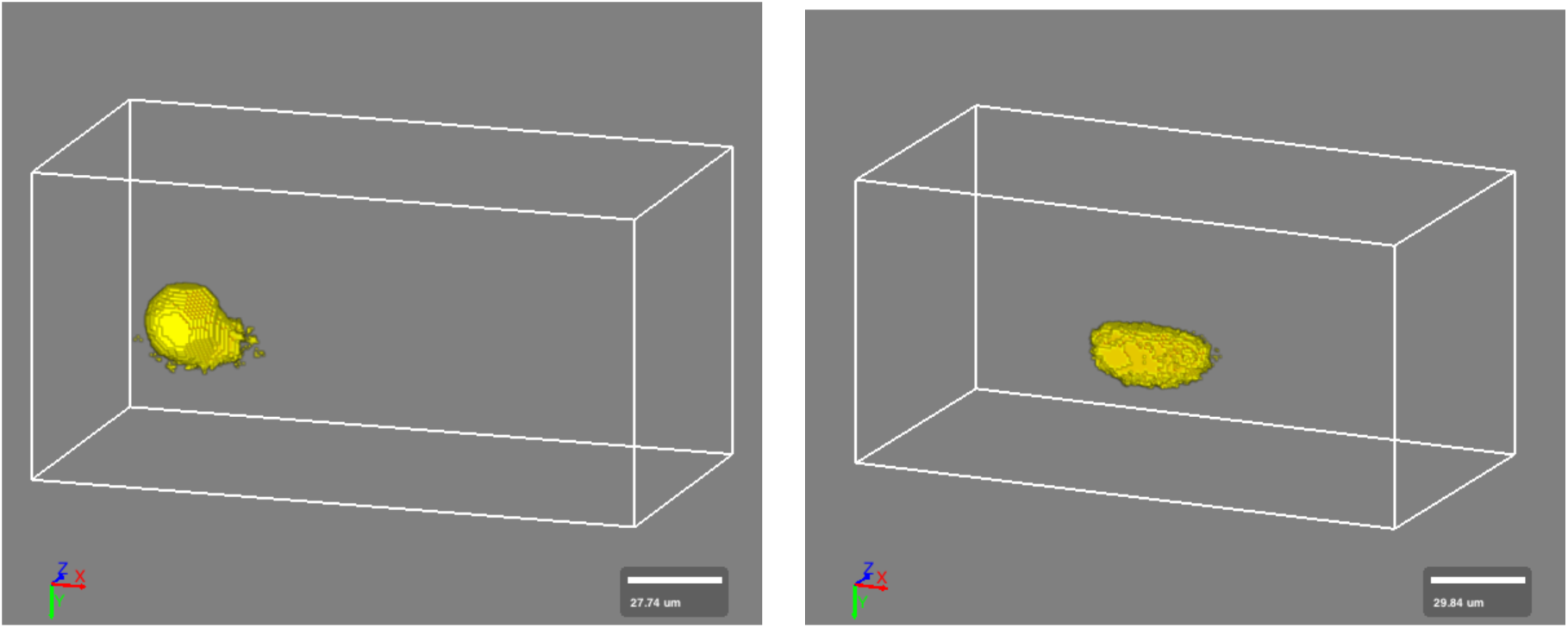
Results of 3D simulations of a single cell moving up gradient of an external field from the left toward the right of a cuboid, taken at 50 MCS (left) and 800 MCS (right). See movie 4 in the supplementary material.

For 3D simulations which were performed using Morpheus, we used its membrane diffusion feature which allows components attached on the cell membrane to diffuse on the surface area. The equation for membrane MT1-MMP [*M*] is modeled using a partial differential equation (PDE), given by

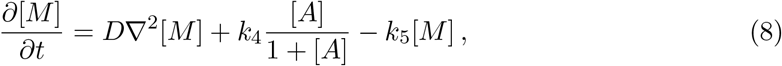

where *D* is the diffusion coefficient, *k*_4_ the production rate due to exocytosis, and *k*_5_ the internalization rate. The PDE contains a 2D diffusion term and two reaction terms. To account for a larger spatial distribution in 3D cells, the equations for inactive proMT1-MMP [*P*] and active MT1-MMP [*A*] and their parameter values are also modified as follows

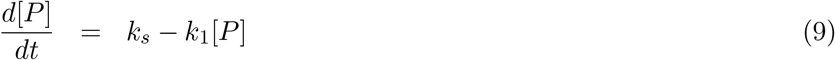

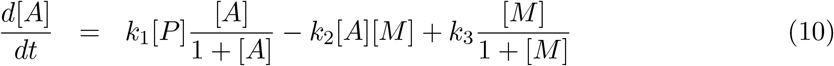

where the parameter values are given in Table 3. All MT1-MMPs are initialized with a similar value, that is [*P*_0_] = [*A*_0_] = [*M*_0_] = 0.5. Periodic boundary conditions are applied on the PDE on the membrane.

**Table 3:** Parameter values for 3D Simulations

| Parameter | Estimated Value |
| --- | --- |
| $k_s$ | 2 |
| $k_1$ | 0.5 |
| $k_2$ | 0.5 |
| $k_3$ | 0.15 |
| $D$ | 0.05 |
| $k_4$ | 0.67 |
| $k_5$ | 0.5 |
| $\phi$ | 80 |

Because membrane MT1-MMP [*M*] diffuses on the cell surface, only a fraction of [*M*] is localized at invadopodia and the amount is given by

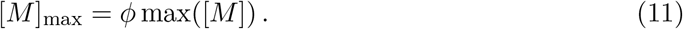

In 3D simulations, the maximum value of [*M*]_max_ in Eq. (11) replaces [*M*] in the membrane Hamiltonian,

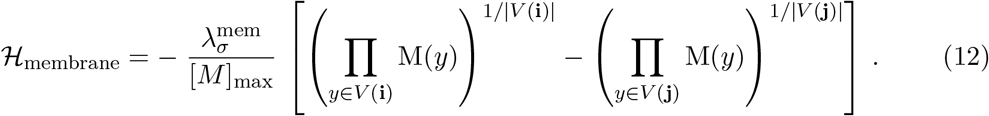

Using all of these modified equations and parameters, we performed 3D simulations with one value of cell-matrix adhesion energy, which is *J*_cell-medium_ = 150 and show the results at 50 MCS (left) and 800 MCS (right) in Fig. 6. Our 3D chemotactic cell moves forward with a lobopodial protrusion-like structure, as seen in the right figure. The localized membrane MT1-MMP on the surface of our 3D cell is visualized in Fig. 7, where the highest concentrations of [*M*]_max_ occur at the front edge, in the direction of cell movement.

**Figure 7:**
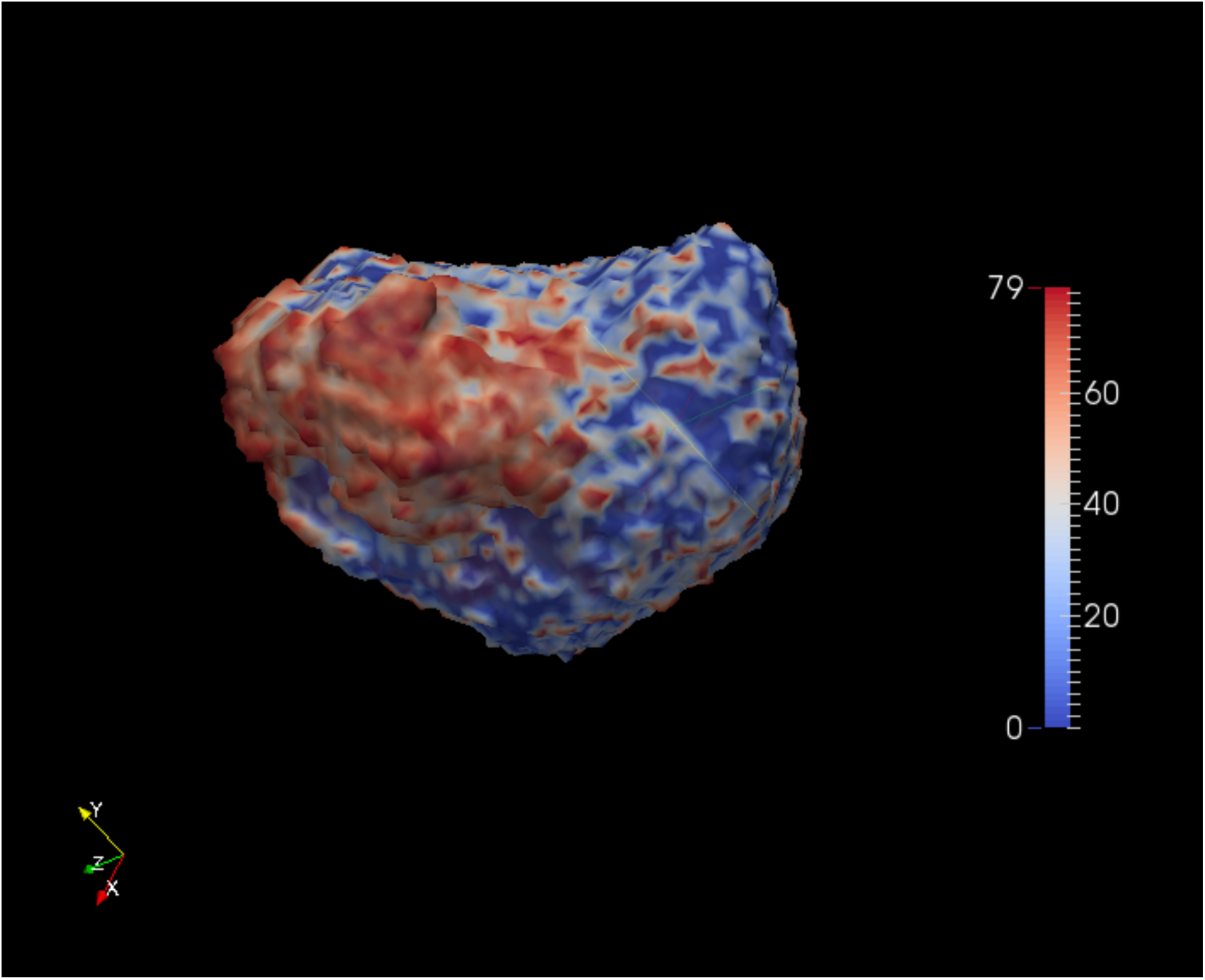
Localization of MT1-MMP on the cell surface in a 3D simulation.

## 4 Discussion and Conclusions

How cells migrate and what factors drive and inhibit cell migration have been intriguing researchers since the last few decades, because, the answers to these questions hold keys to the cure of so many diseases that are impacted by migrating cells. MT1-MMP has long been considered one of the prominent enzymes responsible for promoting cell migration. Aside from its role in the proteolysis of ECM components, MT1-MMP has also been experimentally studied for its role in stimulating cell migration by destabilizing, hence disassembling, strong cell-matrix adhesions. The destabilization is achieved by shedding cell transmembrane receptors from the cell surface which leads to disassembling of stable cell-matrix adhesions and prompting cells to detach from strong adhesive structures that have prevented cells from moving forward.

In this paper, we have developed a multiscale model of single cell migration driven by cell-matrix adhesion destabilization and disassembly by the action of MT1-MMP. In the model, the CPM is used to model cellular behaviors, such as cell motility by enhancing membrane protrusion and retraction, polarization, and motile cell morphology. All of these behaviors are influenced by the intracellular and membrane dynamics of MT1-MMP modeled by differential equations.

Our multiscale model has successfully simulated motile cells with intensified membrane fluctuations caused by MT1-MMP dynamics. Using the model, still cells that are strongly attached to the ECM can weaken their cell-matrix adhesion and eventually migrate. The model also produces cell polarization, in which MT1-MMP on the cell surface is localized to the front edge during migration both in both 2D and 3D simulations, as has been observed in experiments [30, 33, 50]. Our computational simulation results also show the effect of MT1-MMP expression on cell morphology, where very low to no expression of MT1-MMP produces a rounded shape whereas relatively high expression is related to a spreading morphology, as shown in Fig. 2. The expression of MT1-MMP affecting cell morphology has been observed experimentally. MKN45 cells that were transfected with MT1-MMP showed a spread morphology on Laminin-5 [76]. HT-1080 fibrosarcoma and MDA-MB-231 carcinoma cells whose MMP expression, along with serine proteases, cathepsins, and other proteases were inhibited induced a spherical morphology [77]. Elongated cell morphology is particularly visible during directed cell movement or chemotaxis. Simulations of chemotaxis in both 2D and 3D show almost similar elongated cells with a blunt, cylindrical lobopodial protrusion, as seen in Figs. 5 and 6. Cells that have lobopodial protrusion have been experimentally observed to be dependent on cell-matrix adhesion for their motility. Depending on their environment, particularly 3D migrating cells use a wide variety of modes of migration. The environment dictates the amount of cell-matrix adhesion and actomyosin contractility. Lobopodial fibroblasts, for example, require stronger adhesions than rounded amoeboid cells [78]. Polarization, along with chemosensing and locomotion, are steps taken by cells in conducting chemotaxis and these steps have been observed in various types of cancer [79].

Processing cell transmembrane receptors responsible for cell adhesion by MT1-MMP is part of MT1-MMP-dependent cell migration. Our model is able to simulate this role of MT1-MMP in stimulating cell migration. Our model can be used together with models of ECM proteolysis for an integrated representation of MT1-MMP-dependent models of cell migration, or, it can be applied on its own for modeling cell migration that relies on cell-matrix adhesion destabilization and disassembly. The results of computational simulation of our model indicate that targeting the shedding process by MT1-MMP is as important as targeting ECM proteolysis by MT1-MMP in our quest to discover effective and efficient disease treatment and drug discovery. To achieve qualitative and quantitative results that are in line with *in vitro* and *in vivo* experiments, our future work will include data-driven modeling of MT1-MMP and cell transmembrane receptor synthesis and expression using machine learning algorithms.

Despite our understanding of MT1-MMP-mediated shedding of ECM components that plays a significant role in cell migration has broadened our knowledge on MT1-MMP capability, however, experimental studies in this area have not progressed much since the last twenty or so years ago. Detailed mechanisms of shedding down to its molecular level still need to be investigated, because, further insight into the shedding of ECM components may identify more precise target processes for the development of the disease treatment in the future.

